# Genetics of phosphorus use efficiency in a MAGIC wheat population grown in the field

**DOI:** 10.1101/2020.08.27.271205

**Authors:** Anton P. Wasson, Alexander B. Zwart, Arunas P. Verbyla, Gilbert Permalloo, Chandrakumara Weligama, Peter R. Ryan, Emmanuel Delhaize

## Abstract

Phosphorus (P) is an essential plant nutrient and regular applications are essential in most farming systems to maintain high yields. Yet the P fertilizers applied to crops and pastures are derived from non-renewable resources. It is therefore important to find agronomic and genetic strategies for using this resource efficiently, especially since only a proportion of the applied P is absorbed by crops. The aim of this study was to identify Quantitative Trait Loci (QTL) for P use efficiency (PUE) in wheat using a Multiparent Advanced Generation InterCross (MAGIC) population grown in the field. The 357 genotypes were arranged in paired plots with and without P fertilization. Yield and biomass were measured and PUE was calculated as either the performance of the genotype relative to the average response to fertilization, or the performance of the genotype relative to the average resilience in the absence of fertilization. Five trials were conducted over three years in Australia at three sites with contrasting clay and sandy soil types.

Genotypic variation for response and resilience were identified in all trials with moderate to strong correlation with productivity with and without P between trials. Multiparent Whole Genome Average Interval Mapping (WGAIM) QTL analyses were conducted on the four traits (Biomass / Yield × P Response / Resilience) across the five trials and identified 130 QTL in total. QTL within 10 cM of each other were clustered into 56 groups that were likely to represent identical or linked loci. Of the clusters 27 (48%) contained only a single QTL but 17 (31%) contained 3 or more in different trials or traits. This suggests multiple biological mechanisms driving PUE in different environments. Eight of the 56 groups collocated with QTL for seedling root hair length identified in the same MAGIC population in an earlier study.

**Highlight:** Identification of genetic loci for phosphorus use efficiency in a multigenic population of Australian wheats grown on contrasting soils.

## Introduction

Phosphorus (P) is an essential macronutrient critical for many molecules and processes including nucleic acids, phospholipids, ATP, and phosphorylation reactions (Vance *et al*., 2003). P fertilizer is widely applied in Australia where soils are typically low in plant-available P. In 2016/17 2.4 MT of P fertilizers were applied to 23.8 million ha of land in Australia (Australian Bureau of Statistics, 2018). P fertilizers are a non-renewable resource because a large fraction of the world’s supply is derived from a few large deposits of rock phosphate. The input costs associated with the production and delivery of P fertilizers are also rising and increasingly erode the profitability of crop production. The over-use of P fertilizers can lead to significant environmental harm as P enters waterways through soil erosion or leaching (Sharpley *et al*., 2001; Ulén *et al*., 2007). Hence improving the efficiency of P uptake and utilization by crops are important goals for breeders.

Plants absorb P from the soil as soluble inorganic phosphate (Pi), mainly H_2_PO_4_ ^-^ (Bieleski, 1973; Ullrich-Eberius *et al*., 1984; Sakano, 1990; Schachtman *et al*., 1998). However, the concentration of soluble Pi in soils is often very low because Pi is rapidly adsorbed onto mineral surfaces, incorporated into organic compounds or bound by cations (e.g. Fe^2+^, Al^3+^, Ca^2+^) to form poorly soluble complexes. Consequently, Pi is poorly mobile and tends to accumulate in the upper layers of the soil. The absorption of Pi by plants can be limited by its slow rate of diffusion towards the root (Shen *et al*., 2011). Isotopic studies indicate that as little as 9-23% of applied Pi is utilized by wheat in the year of application (Mclaughlin and Alston, 1986; Sharpley, 1986; Mclaughlin *et al*., 1988). These sparingly-available pools of P can be accessed by crops in subsequent years (Simpson *et al*. 2011) especially if plants have mechanisms to increase their solubility or to mine a greater volume of soil.

Traits that improve the efficiency with which plants use Pi relate to Pi uptake, P utilization or P signalling pathways (Bovill *et al*., 2013). Therefore phosphorus-use efficiency (PUE) can be defined in various ways and at various scales depending on the constraints that restrict the uptake of Pi or its conversion into harvestable product. Pi uptake is influenced by root system architecture, symbiotic relationships with arbuscular mycorrhizal fungi, root exudates that improve P mobilisation in the soil, the expression of Pi transporters in the roots and xylem loading. An idiotypic root system architecture for maximizing Pi uptake might include shallow roots to enhance topsoil foraging, adventitious axial roots, more lateral roots and branching, and long root hairs. By contrast, traits that affect utilization enable plants to more efficiently convert the Pi absorbed into biomass and grain by recycling, remobilizing, and translocating the Pi to the most important tissues and prioritising metabolic processes. All these processes are controlled by signalling pathways that function at multiple levels including through gene expression with miRNAs and transcription factors; through protein modification (kinases, ubiquitination); and through sugar signalling.

Consequently, the benefit of one strategy for improving PUE over another strategy depends on the nature of the constraint in each environment. Traits that improve Pi uptake efficiency will be more important for highly Pi-fixing soils (Manske *et al*., 2001), whereas traits that improve Pi utilisation efficiency will be more useful on soils where P availability in not limiting. Genetics and traits that improve PUE in an environment with one type of constraint may be of little benefit in another environment with a different constraint. Genetics and traits may even counteract each other since uptake and utilization can sometimes be negatively correlated (Wissuwa *et al*., 1998; Su *et al*., 2009; Rose *et al*., 2011).

Varietal differences in the efficiency of nutrient uptake have interested researchers for many years, with studies on the “phosphorus feeding capacity” of maize genotypes appearing as early as 1936 (Lyness, 1936). Bovill *et al*. (2013) suggested that breeding for yield potential under high P fertilizer rates would passively increase PUE and there is some evidence that this occurs (Batten, 1992; Ortiz-Monasterio R. *et al*., 1997; Egle *et al*., 1999). Nevertheless, it is likely that PUE can be improved still further by targeted breeding for specific traits.

Most genetic studies of PUE in wheat have used biparental populations in pot trials or hydroponic experiments and relatively few incorporated full-season field trials. Furthermore, most used winter wheats and low-density genetic maps. Nevertheless, these reports provide a useful basis for further work.

Su *et al*. (2006) identified 39 QTL for “P deficiency tolerance” in a pot trial by screening a doubled haploid (DH) population generated from Lovrin10 and Chinese Spring parents. Three major QTL clusters were identified on chromosomes 4B, 5A, and 5D, the latter two being linked with the vernalisation genes *Vrn-A1* and *Vrn-D1*. The winter alleles of *VRN1* genes on 5A, 5B, and 5D were later implicated in shallower root angles in wheat (Voss-Fels *et al*., 2018). Su *et al*. (2009) subsequently performed pot experiments and field trials using DH lines from winter wheat parents Hanxuan 10 and Lumai 14. They measured P uptake and biomass in different P treatments and detected a total of 195 QTL, seven of which were strongly linked with uptake efficiency and six with utilization efficiency. P uptake efficiency tended to be negatively correlated with utilisation efficiency in that study and only two loci on 3A and 3B positively influenced both traits.

Guo *et al*. (2012) used hydroponic experiments to estimate nutrient-use efficiency for N, P, and K in a set of recombinant inbred lines (RILs) of winter wheat (Chuan 35050 and Shannong 483). Of the 380 QTLs detected almost half co-located in 10 clusters some of which were associated with improved uptake and utilisation of all three nutrients. Zhang and Wang (2015) also used hydroponics to score three sets of RILs for a range of traits. They identified 110 QTL with 28 of the major QTL falling into 18 clusters. Yuan *et al*. (2017) studied a range of traits on seedlings and mature plants in 184 RILs (TN18/LM6) in hydroponic screens and field trials at low and high P. A total of 163 QTL were identified, many of which co-located into 10 clusters on chromosomes 1A, 1D, 4B, 5D, 6A, and 6B. PUE was strongly correlated with various measures of biomass and yield in seedlings and mature plants and the authors concluded that some simple morphological indexes could be used by breeders to evaluate PUE on a large-scale.

Ryan *et al*. (2015) screened two biparental populations for biomass in a series of glasshouse experiments using a highly P-fixing soil. Seven significant QTL were identified from a set of RILs (Chuan Mai 18/ Vigour 18), with the largest on 7A, and nine QTL were detected in a DH population (Kukri/Janz) with two located on chromosomes 4B and 4D (likely *Rht-B1a* and *Rht-D1a*) accounting for 25% of the total variance. The authors concluded that early vigour contributed positively to PUE in both populations.

Yang *et al*. (2021) screened a DH population, derived from Yangmai 16 and Zhongmei 895, for seedling root and biomass traits in hydroponics in zero, low, and high P conditions. Using a ∼10 k marker linkage map they identified 34 QTL in 7 clusters with pleiotropic effects on traits including root length, the number of root tips, and root surface area.

The aim of the present study was to identify QTL for ‘P responsiveness’ and ‘P resilience’ in wheat, both of which are independent of the standard measures of productivity, biomass and grain yield. The benefit of being independent of absolute productivity at a given P level is that any QTL identified are likely to be specific for PUE. Explanations for these traits are provided later.

We used a Multiparent Advanced Generation InterCross (MAGIC) population constructed from four successful Australian cultivars from different regions of the Australian wheatbelt (Huang *et al*., 2012, 2013). The MAGIC methodology enables the identification of many small effect QTL and is well-suited for studying PUE at multiple sites across Australia. The same population has previously been used to identify QTL for hair length (Delhaize *et al*., 2015), coleoptile and seedling growth (Rebetzke *et al*., 2014), paired spikelet formation (Boden *et al*., 2015), grain dormancy (Barrero *et al*., 2015), and canopy architecture (Richards *et al*., 2019). Some of these traits may also affect nutrient efficiency. For instance, long root hairs have been identified as a PUE trait in controlled environment studies (Gahoonia and Nielsen, 1997, 2003), and simulations suggest root hairs may be responsible for 50% of plant P uptake (Ruiz *et al*., 2020).

## Methods

### Plant material

The MAGIC population was constructed from four Australian wheat cultivars (Baxter, Chara, Westonia, and Yitpi) grown in different regions of the Australian wheatbelt. Used in conjunction with a high-density genetic map and a 90K single nucleotide polymorphism (SNP) chip (Cavanagh *et al*., 2013; Wang *et al*., 2014) it is specifically designed for mapping many small-effect QTL (Huang *et al*., 2012, 2013). Our trials used a subset of the population comprised of 357 semi-dwarf genotypes (Huang *et al*., 2013) which were chosen to maximise allelic diversity. All genotypes in the population carried the *Rht-B1* or *Rht-D1* alleles and excluded tall and double-dwarf genotypes. Another group of wheat cultivars were included in the trails (usually 11 to 17) to calibrate the measurements of biomass between quadrat sampling and Light Detection and Ranging (LIDAR) instruments.

### Field Trials

Trials were conducted on farmers’ fields at Ardlethan in New South Wales (NSW), Wallaroo in the Australian Capital Territory (ACT), and York in Western Australia (WA) (Table 1). The sites were identified as being P responsive for wheat growth as determined with a bioassay, with plant available P measured using the Colwell P assay (Colwell, 1963). For the bioassay rows of wheat seedlings (cv. Scepter) were sown into a tray of topsoil (0-10 cm depth) with and without triple superphosphate (19% P; 100 mg kg^-1^ P where applied). They were grown in the glasshouse for 10 days and then seedling size was assessed to establish if there was a response to P fertilization.

**Table 1.** Site Details

|  | Ardlethan (NSW) |  | Walleroo (ACT) | York (WA) |  |
| --- | --- | --- | --- | --- | --- |
| **Soil type** | Clay |  | Clay | Sand over Clay/Loam |  |
| **Phosphate Buffering Index (PBI)** | 71 (Low) |  | 50 (Very low) | 28 (Very very low) |  |
| **Sowing Configuration** | 6 m plots (5 m harvested). 100 plants/m<sup>2</sup> target. 6 rows, 0.33 m spacing, 2 m centres. |  | 4 m plots (3 harvested). 180 plants/m<sup>2</sup> target. 10 rows, 0.18 m spacing, 2.2 m centres. | 6 m plots (4 m harvested). 120 plants/m<sup>2</sup> target. 6 rows, 0.254 m spacing, 1.75 m centres. |  |
|  | 2017 | 2018 | 2018 | 2017 | 2018 |
| **Latitude** | -34.142081° | -34.134528° | -35.174990° | -31.939802° | -31.940146° |
| **Longitude** | 146.854074° | 146.855161° | 149.044366° | 116.898947° | 116.903877° |
| **Sowing Date** | 2017-05-23 | 2018-06-05 | 2018-06-13 | 2017-06-06 | 2018-06-08 |
| **Harvest Date** | 2017-11-23 | 2018-12-01 | 2018-12-19 | 2018-12-14 | 2018-12-17 |
| **pH** | 4.6 | 5.3 | 4.7 | 5.1 | 4.6 |
| **Colwell P (0-10 cm)** | 13.2 | 24.8 | 12.0 | 25.7 | 30.0 |

In the field, genotypes were sown in paired plots with and without the P fertilizer treatment (0 and 30 kg P ha^-1^ respectively), applied as triple superphosphate which was added with the seed in runs of the plot seeder at sowing. The two P treatments were randomised to the runs within each successive pair of runs across the trial. Genotypes were randomised to the plots within each pair of plots formed by the intersection of ‘run pairs’ with ranges. The full set of genotype × treatment combinations were replicated across two blocks in each trial. The optimal randomisations for these designs were produced using the ‘od’ statistical software package (Butler 2019; Butler 2013) in the R statistical computing environment.

Further details of the trials are found in Table 1. Nitrogen was supplied as urea prior to sowing and was top-dressed again at stem elongation with urea (timed with rainfall), or liquid N fertilizers (e.g. urea and ammonium nitrate liquid formulations - UAN). Muriate and sulphate of potash (KCl and K_2_SO_4_) were applied as required to ensure adequate K and S, as were micronutrients. Foliar diseases were managed with prophylactic fungicide and pesticide applications. Weeds were managed with district herbicide application practices at the recommended rates.

The trial at York in 2017 was affected by *Rhizoctonia*. Damage to the plots was scored and the data used as a covariate in the spatial model of the trial to remove the effect on productivity. Affected parts of the plots were excluded from the measurement of biomass.

Temperature and rainfall are summarised in Figure 1. Higher than average temperatures were experienced during grain filling for all trials except York 2018. A colder than average start was experienced at Ardlethan in 2017. The Ardlethan and Wallaroo trials experienced much lower than average in-season rainfall (rain events at the end of the season in Ardlethan 2017 and Wallaroo coming too late to contribute to productivity) whereas York experienced average rainfall.

**Figure 1.**
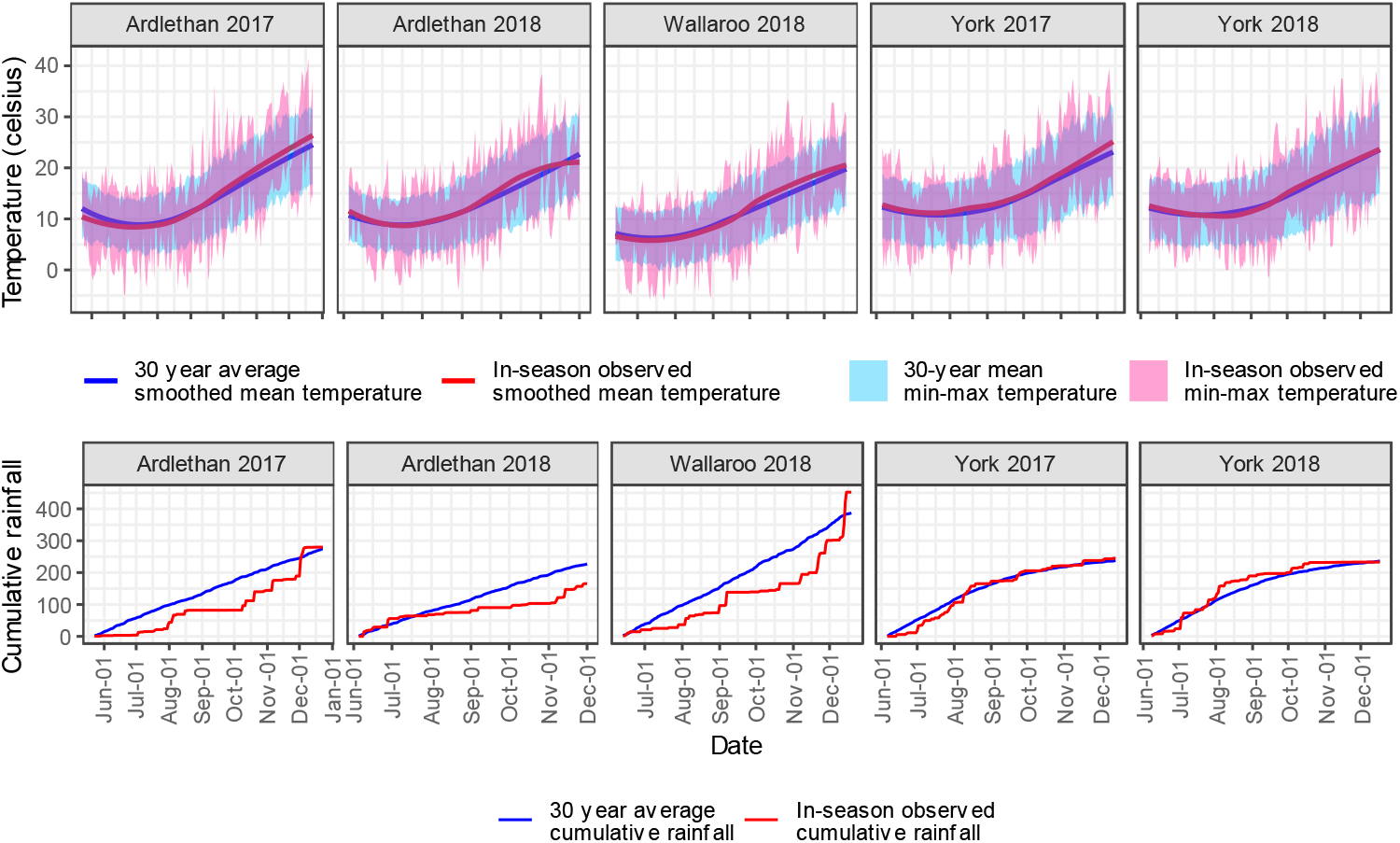
In-season weather relative to the climatic average. The top row shows the temperature and the bottom row shows the cumulative rainfall. The 30-year average (to 2017) is shown in blue, and the in-season observations are shown in pink. For temperature, the ribbon is the average daily maximum and minimum temperature, with the line the daily average with a loess smoother.

### Phenotyping

Biomass was measured using a terrestrial LIDAR phenotyping system, the Phenomobile-Lite, using a 3D voxel index methodology (Jimenez-Berni *et al*., 2018). Four inner rows were sampled by quadrat from a subset of plots (between 44 and 84 plots were sampled per trial). Quadrat sizes varied from 0.21 to 0.66 m^2^. A model of 301 quadrat cut biomass measurements (n=44–84 in each trial) and LIDAR biomass measurements was generated with a multiple R^2^ of 0.88; indicating that the two measurements were comparable. However, the LIDAR measurements integrated a larger area of the plot, were less susceptible to handling losses, and were faster to perform; factors that made LIDAR the method of choice.

Yield was measured with a trial plot header.

### Statistical analysis and QTL mapping

Analysis methodology followed that described in McDonald *et al*. (2015). First, for each productivity measure (yield or biomass) a Multi-Environment Trial (MET) analysis of the combined data of the 5 trials was conducted using a linear mixed model formulation. The analyses accounted for the blocking and unit structure in the design of each field trial through the inclusion of corresponding random effects. Spatial effects such as large length scale trends or random run/range effects could also be accounted for by including appropriate random or fixed effects, and the presence of autocorrelated error structure across run/ranges could also be modelled. The impact of *Rhizoctonia* infection in the York 2017 trial was accounted for by the addition of fixed linear trend and curvilinear spline components, as per Verbyla *et al*. (1999).

Genetic effects were captured by the inclusion of a genotype by P treatment by environment term in the model, allowing the estimation of the genetic variance-covariance matrix across the treatment-environment combinations (as there are 5 trials and 2 P treatments, a 10 by 10 matrix) and the associated genetic effect Best Linear Unbiased Predictors (BLUPs). Since estimation of the full unstructured genetic variance matrix can be computationally difficult (55 parameters to be estimated), it was instead approximated using the factor-analytic approach proposed by Smith *et al*. (2001, see also 2005).

P response in each trial is defined using the conditional distribution of the true ‘with P’ genetic effect given the ‘without P’ genetic effect for each trial. Under the normality assumption in the linear mixed model, the mean of this distribution is a linear regression of the ‘with P’ genetic effect on the ‘without P’ genetic effect (with zero intercept), with the regression slope parameter being the ratio of the genetic covariance between the two treatments to the variance of the ‘without P’ treatment genetic effect. Under this model, the residual is then the difference between the ‘with P’ genetic effect and the regression coefficient multiplied by the ‘without P’ genetic effect. The estimated residual is found by replacing the genetic effects by their BLUPs, and the covariance and variance in the regression coefficient by their estimates found in fitting the linear mixed model. The estimated residual is a measure of the P response.

The regression slope for ‘with P’ response represents the genetic productivity increase per unit increase in the genetic effect under the ‘without P’ fertilization. Thus, the residual for a genotype is the difference in the ‘with P’ genetic effect from the appropriately adjusted ‘without P’ genetic effect. A large positive predicted residual indicates that the genotype responds strongly to added P. The concept is illustrated in Figure 2.

**Figure 2.**
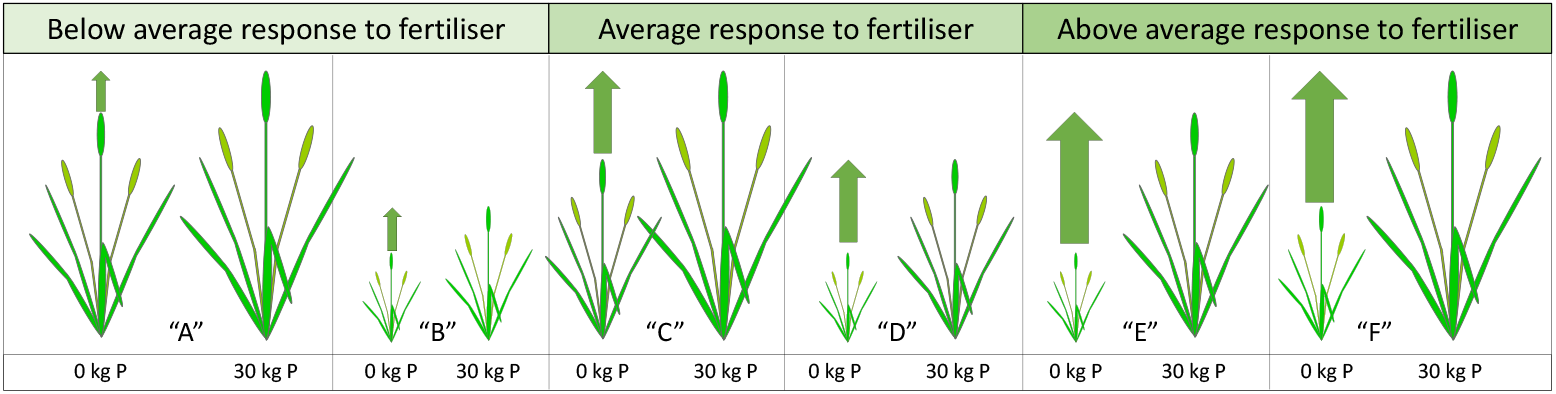
Cartoon illustrating how genotypic “response” to P fertilisation might not relate to overall productivity. Pairs “A” and “B” are less responsive, (“C” and “D” show an average response,) and “E” and “F” are highly responsive. The “resilience” can be thought of as the inverse of this change: i.e. how much does the productivity of a genotype decrease from a fertilized to an unfertilised state. Consequently, “A” and “B” are more resilient to the low P treatment than “E” and “F” which show low resilience to the low P treatment. Note that while the magnitude of the actual change for a given genotype is the same for resilience and response, how that change compares to the **population** average response or resilience (i.e. the residual upon which the metric is based) will be different because it is calculated relative to different axes. The only circumstances under which they would be the same is if the genetic variance of productivity at both P levels were equal.

This study introduces a second criterion, “P resilience”. The well-fertilized productivity is taken to be the norm (true in the case of conventional yield breeding for industrial agriculture) and we consider the loss in productivity under low P conditions, relative to the population average. Resilience represents the genetic productivity loss per unit decrease in the ‘without P’ genetic effect. In this case a positive residual indicates a higher than average productivity without P fertilization, i.e. greater resilience.

In this paper, P resilience is defined similarly to response, by reversing the roles of the ‘with P’ and ‘without P’ treatment genetic effects. Conceptually, resilience appears to be the inverse of response, but mathematically it is not because the residual is expressed in the units of the dependent variable and will hence be different. The same covariance is used in the regression coefficient in both cases, but for response it is divided by the variance of the ‘without P’ treatment, while for resilience it is divided by the variance of the ‘with P’ treatment (Figure 2).

This approach can be contrasted with an alternative such as relative yield (e.g. Gong *et al*., 2016). Relative yield is simply the yield at low P expressed as a percentage of yield at high P. However, if every genotype has the same fixed yield response to fertilization then those genotypes with a smaller absolute yield with have a larger percentage decline, or smaller relative yield, than those with higher absolute yield. This means that relative yield is strongly reflective of absolute yield.

Finally, each of the resulting 20 datasets (yield/biomass × resilience/response for each of five trials) were searched for QTL using the mpwgaim R package (Verbyla *et al*., 2014).

The above analyses were conducted in the R (version 3.5.1) statistical computing environment (R Core Team, 2020), with the asreml package (version 3.1) for R used for all linear mixed model fits (Butler *et al*., 2009).

## Results

Five field trials were performed at three sites with P-responsive soils over two years: Ardlethan in 2017 and 2018, Wallaroo in 2018, and York in 2017 and 2018. The two measures of productivity calculated from the paired plots were anthesis biomass and grain yield. Measurements made on each plot were subject to a spatial analysis using mixed models that removed site trends from the data. The resulting measures are residuals that represent the deviation from the trial average genetic change in productivity between fertilization states of a measured genotype at a given site.

### Effect of P treatment on productivity

Figure 3 shows the distributions of productivity across the five trials at the two P fertilization levels. As expected, fertilisation with P increased both yield and biomass in all trials (Figure 3) indicating that P was a limiting factor and that the trials were appropriate for the assessment of PUE.

**Figure 3.**
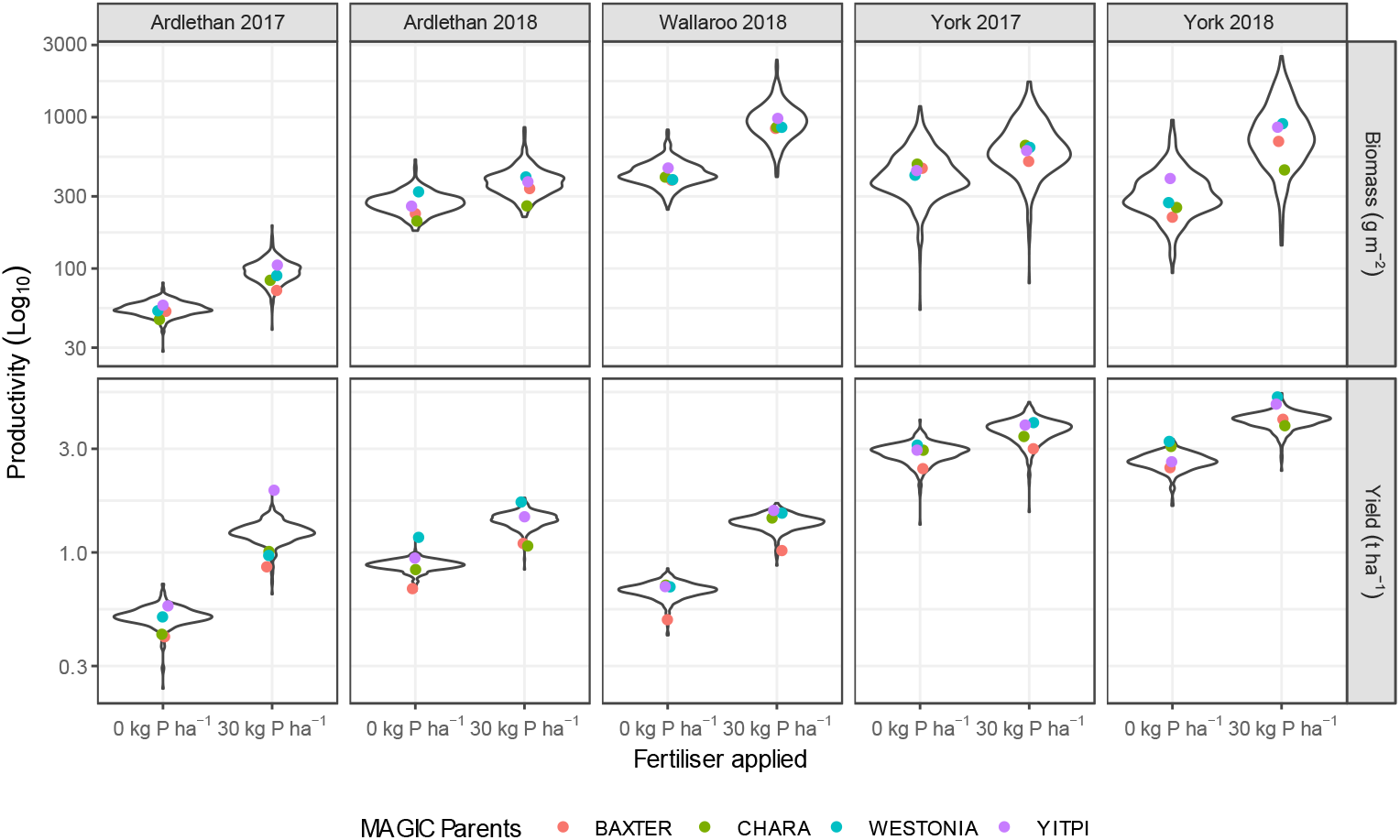
Effect of P on productivity across trials. The plot is faceted by trial from left to right, and by the productivity measures of biomass (top) and yield (bottom). Within each facet the fertilization treatment is on the x-axis and the measure of productivity is shown on a Log10 transformed y-axis. The spread of values across the population is shown in a “violin plot”, the width of the enclosed area reflects the density of the data located there. The logarithmically scaled y-axis shows the distribution within the trial, while allowing a meaningful comparison of the differences between trials. The performance of the cultivar parents of the 4-way MAGIC population, as a comparison to the overall population, is shown by way of coloured points. Productivity uses adjusted BLUPs, being the trial × P fertilizer applied mean plus the genetic response of the genotype. Furthermore, the biomass has been converted from LIDAR index to g m^-2^ using the modelled relationship (see Methods).

The in-season rainfall and temperature (Figure 1) at the sites strongly influenced productivity: York was more productive than the other sites for both biomass and yield whereas Wallaroo was slightly more productive for biomass than Ardlethan (Figure 3). The population extremes for productivity typically exceeded the range of variation of the four parental cultivars, particularly for biomass (Figure 3). Among the parents cv. Yitpi typically had the greatest biomass, while cv. Westonia yielded better at York and Ardlethan in 2018 (Figure 3). Cultivars Chara and Baxter typically had the lowest biomass and Baxter was the lowest yielding of the parents (Figure 3).

### Productivity, Response and Resilience across environments

PUE was assessed from the biomass and yield measurements in two ways: as “response” to fertilization, and “resilience” to the absence of fertilization. Consequently, the four traits measured were biomass resilience, biomass response, yield resilience and yield response. The spatial trends in the underlying productivity data across the five trials were modelled with two separate multisite mixed models for yield and biomass, leaving the genetic trends and unexplained variation (error).

There was a strong linear relationship between productivity with and without fertilisation (Figure 4, Supplementary Figure 1). The meaning of P response and resilience, conceptually introduced in Figure 2, is illustrated in Figure 4 using the genotype BLUPs for yield at York in 2018. The 30 kg P ha^-1^ BLUPs are plotted against those at 0 kg P ha^-1^. The BLUPs represent the genetic component of the variation in yield in that trial and fertilization level expressed relative to the population average yield; hence they can be positive or negative. While there is a clear relationship between productivity with the two fertilizer treatments across the whole population (Supplementary Figure 1), of interest are those genotypes that deviate from the relationship; hence those with large residuals, positive and negative, in the regression.

**Figure 4.**
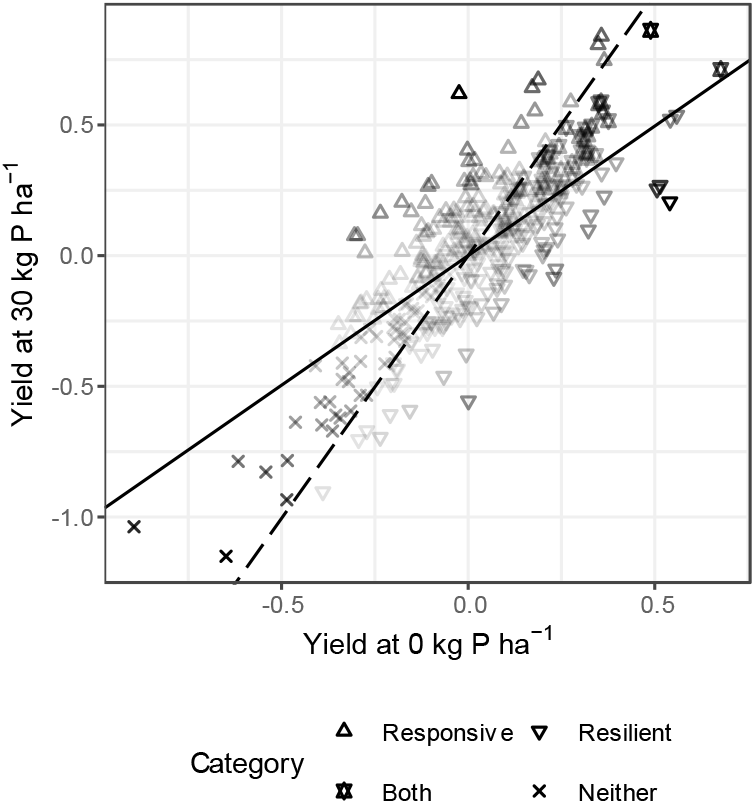
Scatterplot of the genetic component of yield (BLUP) for each genotype at York in 2018. Yield at 30 kg P ha^-1^ is plotted against yield at 0 kg P ha^-1^. Lines for the average genetic yield response to fertilisation (**solid**) and average genetic yield resilience in the absence of fertilization (**dashed**) in this trial. The difference in response for each genotype from the population average is its vertical distance (y-axis) from the average, i.e. the solid line. The difference in resilience for each genotype from the population average is its horizontal distance (x-axis) from the average, i.e. the dashed line. Each genotype has been categorised and coded by shape. If it is **responsive**, i.e. **above** the solid line, it is a **triangle**; if it is **resilient**, i.e. **right** of the dashed line, it is an **inverted triangle**; if it is **both** resilient and responsive, i.e. above the solid line and right of the dashed line, it is a **star**; and if it is **neither** resilient nor responsive, i.e. below the solid line and left of the dashed line, it is a **cross**. The shading of the point indicates the degree of response and/or resilience, and highlights that the measure is independent of yield at either P level. For the genotypes that are neither or both responsive and resilient the shading is the average of the two measures and will increase with distance from the origin. The slope of the regression is derived from a factor analytic model fitted across all five trials in the study and represents the genetic component of the response; it does <u>not</u> reflect a regression of the BLUPs for the two P fertilization levels.

It is important to note that the two regression slopes for response and resilience in Figure 4 are based on the genetic component of yield across all five trials and would differ from a linear regression performed on the BLUPs for any single trial. For response, the genotypes with a large positive residual had an above average response to fertilization with P and those with a large negative residual had a below average response to fertilization with P. Note that these residuals do not reflect absolute yield. The genotypes shaded in Figure 4 as having the highest response to P fertilization are not necessarily the genotypes with the highest yield with P; similarly those shaded with the highest resilience are not those with the highest yield without P. P-resilience is calculated from the inverse relationship; i.e. productivity at 0 kg P ha^-1^ when regressed against 30 kg P ha^-1^ and similar calculations can be made to identify the genotypes showing better and worse resilience compared to the population (Supplementary Figure 1b). Biomass and yield were positively correlated for both response and resilience as discussed below.

It is also important to note that the difference in regression slopes between response and resilience (Figure 4) creates “wedges” where a genotype could be both, or neither, responsive and resilient, but that these genotypes will be strongly correlated with performance at both P levels.

The calculated PUE values across trials for response and resilience are shown in Figure 5. A wider range of distributions were found for response compared to resilience. This reflects the axes on which the trait is expressed: response is expressed on the axis of “productivity at 30 kg P ha^-1^”, which is larger and has more variation than the axis of “productivity at 0 kg P ha^-1^” on which resilience is expressed. Similarly, the variation in yield and biomass were greatest at York which had the greatest productivity over both years.

**Figure 5.**
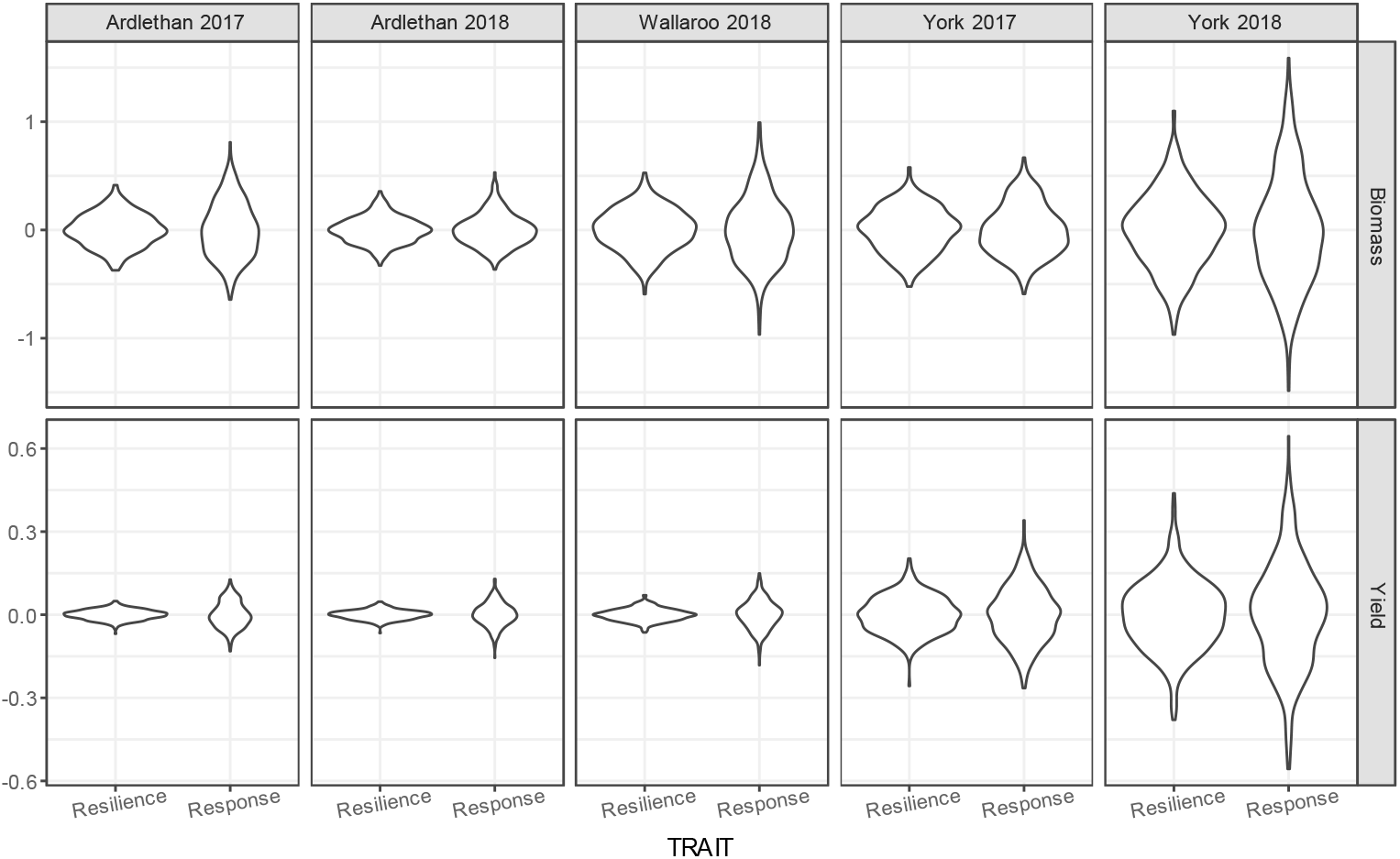
Genotypic variation in PUE across trials. The plot is faceted by trial from left to right, and by productivity (biomass and yield) from top to bottom. Within each facet the PUE trait is on the x-axis and the trait value is shown on the y-axis. Note the variable y-axis is scaled to the extent of variation expressed in each trait.

As noted earlier, response and resilience are distinct from productivity with and without fertilization, respectively, but they were inversely correlated as shown in Figure 6 (a full correlogram appears in Supplementary Figure 2). The strength of the correlations varied with sites and years, but the relationships were generally stronger for biomass than for yield. The strongest correlations occurred at York in 2017 for both biomass (−0.94) and yield (−0.86) but reduced the following year to -0.82 and -0.54, respectively.

**Figure 6.**
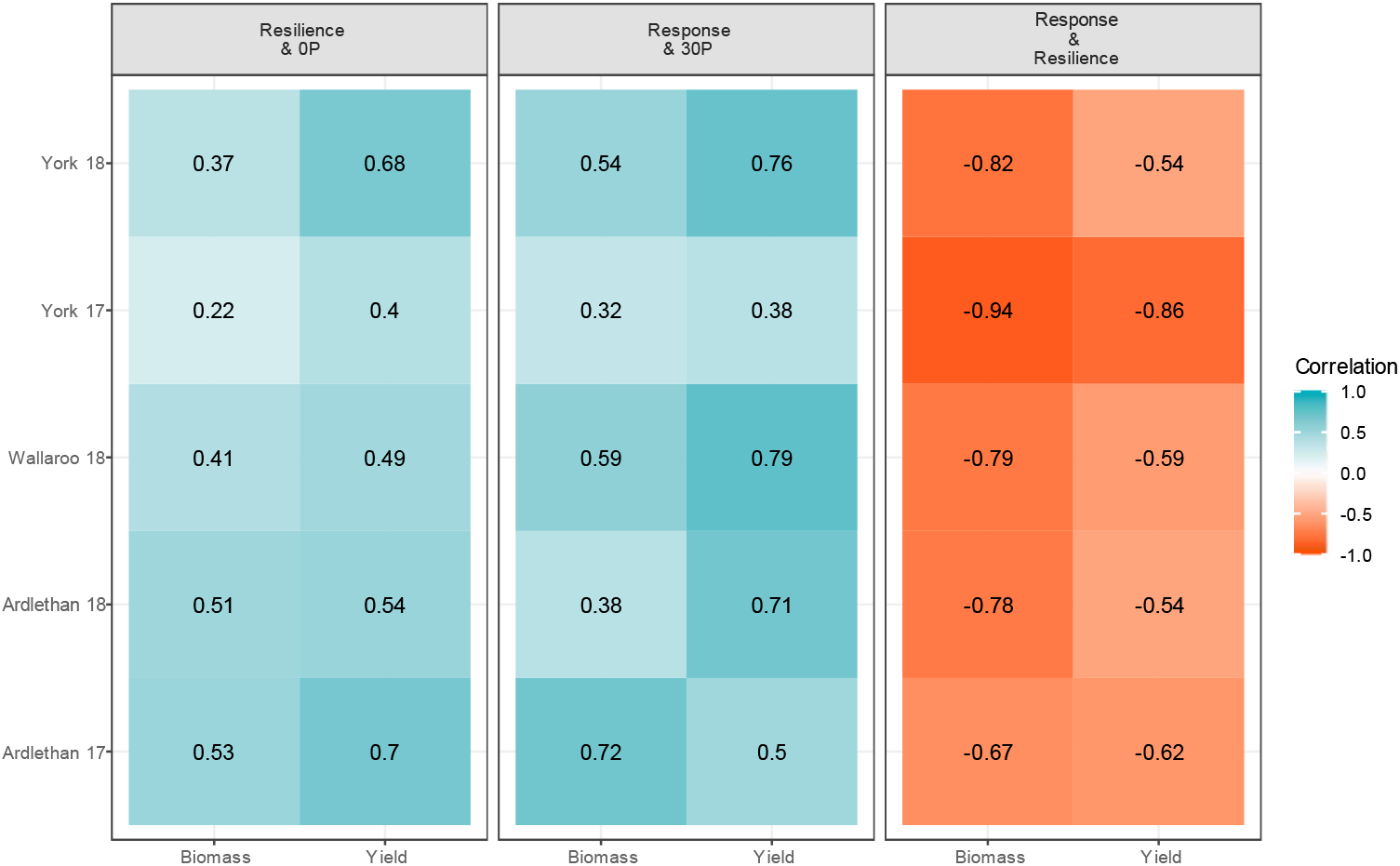
A heatmap of selected correlations between measures. The plot is faceted by the relationship tested: Resilience and Productivity at 0 kg P ha-1, Response and Productivity at 30 kg P ha-1, and Response and Resilience. Within each facet, the productivity measured is on the x-axis, and the trial is on the y-axis. Each tile is coloured and labelled by the size and direction of the correlation (Pearson correlation coefficient).

Resilience is correlated with productivity without fertilization. This relationship was strongest for yield at York in 2018 and Ardlethan in 2017 and weakest at York in 2017. Resilience was only weakly correlated with productivity with fertilization (Supplementary Figure 2).

A similar relationship emerged between response and productivity with fertilization. There was a strong correlation between response and yield at Wallaroo in 2018 and weaker correlations at York and Ardlethan in 2017.

However, there were moderate, negative relationships between biomass resilience and yield response and yield resilience and biomass response in the two York trials and Ardlethan in 2018 (−0.27 to -0.48), but, with the exception of York in 2018, it was not based in an equivalent negative correlation in biomass and yield at either fertilization level.

The heritability of the PUE traits ranged from 0.06 to 0.50 with a mean of 0.26 (Table 2). The lowest heritabilities for yield were obtained for the trials at York in 2017, and the highest for the trials at York in 2018. In 2017 York experienced an early season rainfall deficit, and the trial was affected by a *Rhizoctonia* outbreak. Damage was assessed on individual plots and although accounted for in the spatial modelling, it is possible that this reduced the heritability of the PUE traits. The difference in heritability between resilience and response was less than 0.001 and was rounded out (Table 2). Over all trials, yield was more heritable than biomass (0.32 vs. 0.20).

**Table 2.** Heritabilities of PUE traits across trials.

| Trial | Biomass Response | Biomass Resilience | Yield Response | Yield Resilience |
| --- | --- | --- | --- | --- |
| **Ardlethan 2017** | 0.36 | 0.36 | 0.27 | 0.27 |
| **Ardlethan 2018** | 0.18 | 0.18 | 0.33 | 0.33 |
| **Wallaroo 2018** | 0.25 | 0.25 | 0.41 | 0.41 |
| **York 2017** | 0.06 | 0.06 | 0.11 | 0.11 |
| **York 2018** | 0.17 | 0.17 | 0.5 | 0.5 |

### QTL discovery

Twenty univariate models were generated for the five trials and four traits and the resulting QTL were mapped with MPWGAIM. A total of 130 QTL were identified (Table 3, Supplementary Table 1). QTL were detected on all chromosomes (Figure 7). The most QTL identified for a single trait in a single trial was 14 QTL for biomass response at Ardlethan in 2017. No QTL were detected for yield resilience at Ardlethan in 2018. For the remaining trials the total variance of each trait explained by the QTL ranged from 12.8% (yield response, Ardlethan and York 2018) to 42% (biomass response, Ardlethan 2018) with an overall average of 25.6% (Table 3).

**Table 3.** QTL detected and total variance explained by them (in brackets) for each trial and trait combination.

| Trial | Biomass resilience | Biomass response | Yield resilience | Yield response |
| --- | --- | --- | --- | --- |
| **Ardlethan 2017** | 5 (23.1%) | 14 (42%) | 7 (33.7%) | 7 (20.1%) |
| **Ardlethan 2018** | 9 (27.8%) | 9 (32.1%) | 0 (0%) | 2 (12.8%) |
| **Wallaroo 2018** | 7 (26.9%) | 11 (36.7%) | 4 (15.2%) | 11 (40.2%) |
| **York 2017** | 7 (27.8%) | 7 (27.2%) | 4 (16%) | 4 (17.9%) |
| **York 2018** | 9 (31.3%) | 6 (26.6%) | 5 (16.5%) | 2 (12.8%) |

**Figure 7.**
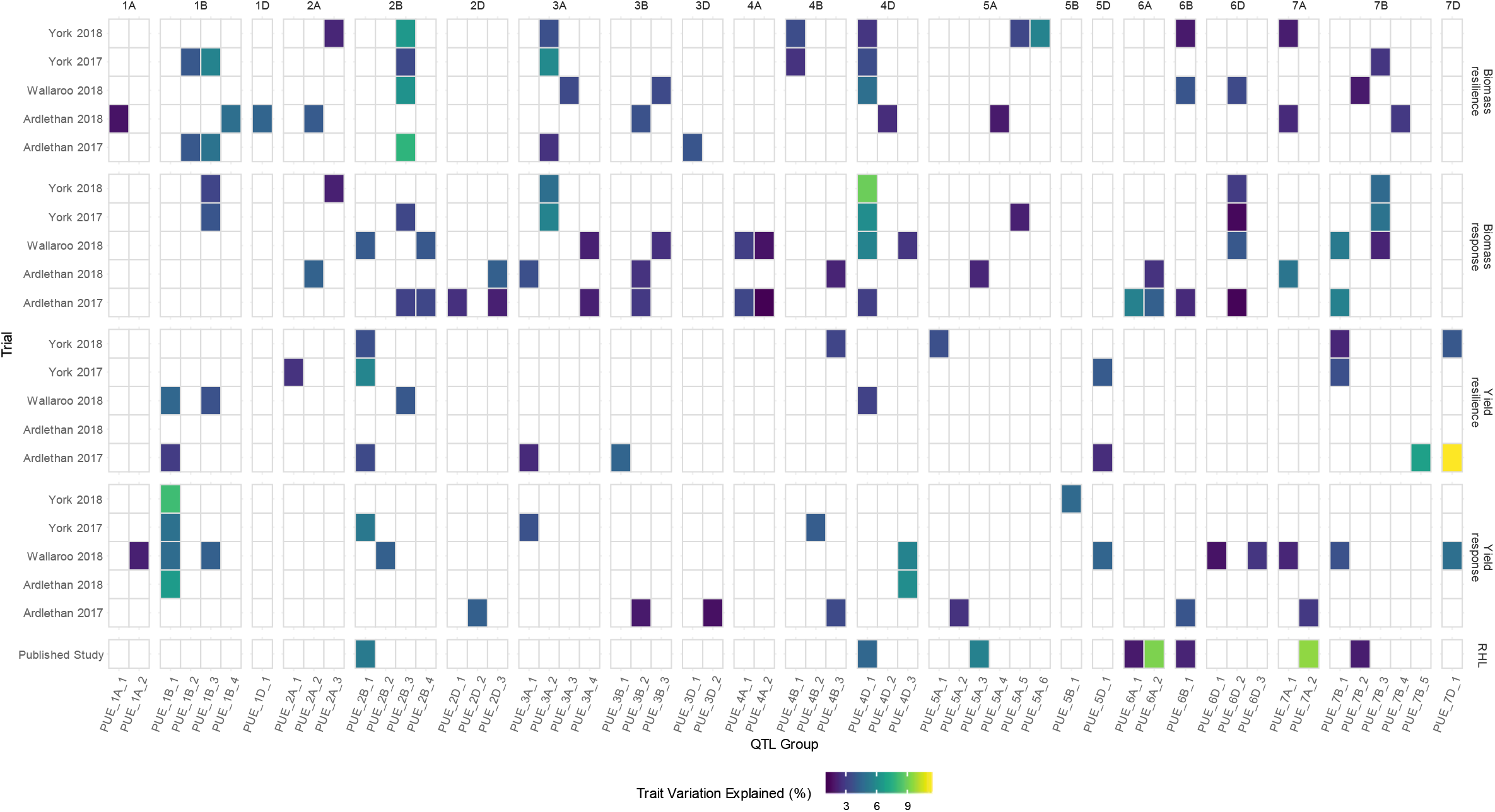
Location (chromosome), occurrence (trial and trait) and strength (variation explained) of the 130 QTL identified for four PUE traits across five trials. The QTL were allocated to 56 groups each designated with a “PUE_” prefix, then grouped and numbered by chromosome. The plot is faceted by chromosome from left to right, and trait from top to bottom. Within each facet the QTL group is on the x-axis and the trial is on the y-axis. The percentage of the genetic variation for the trait explained by the QTL is shown by the colour of the tile. The bottom facet is also a trait, root hair length, from the published data of Delhaize et al. (2015), also from the 4-Way MAGIC population for the purposes of showing co-location.

For breeding programs, the most useful genes are those that positively influence more than one trait over different trials. Furthermore, the most desirable PUE ideotype (Donald, 1968) would efficiently exploit P fertilizer when it was present and soil P when it was not; i.e. it would be both responsive and resilient. However, given the inverse correlation between response and resilience, we were concerned to identify those QTL associated with antagonistic pleiotropy. We considered instances where there were examples of pleiotropy associated with our QTL; i.e. where a QTL for a response at a trial has a collocated (in the same cluster) QTL for resilience. We only considered pleiotropy within the same productivity measure (i.e. biomass or yield) and trial to limit complexity. Where these pleiotropic QTL have founder effects that mirror each other, we can assume it represents a gene driving a mechanism that is antagonistic between response and resilience.

Therefore, we further grouped the 130 QTL into 56 clusters that were within 10 cM of each other on the genetic map as shown in Figure 7. The multiparent WGAIM methodology gives effect sizes for each of the four founders of the population, cv. Yitpi, Baxter, Chara, and Westonia, shown in Figure 8. Of the 56 QTL clusters 27 (48%) only contain a single QTL, and a further 12 (21%) contain only two. The largest cluster contains eight QTL.

**Figure 8.**
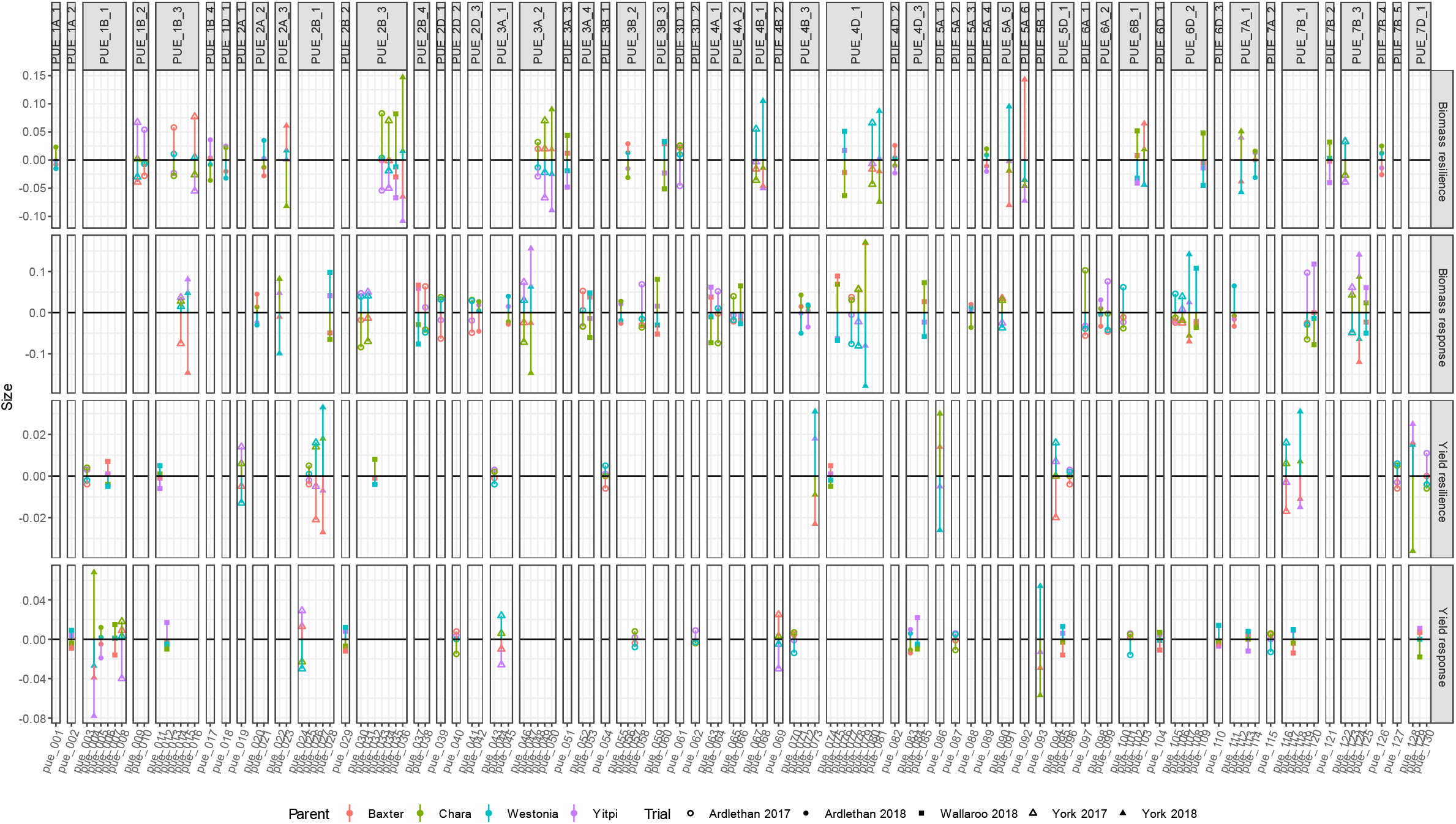
Founder (allele) effects for each of the 130 QTL identified for four PUE traits across five trials in 56 groups each designated with a “PUE_” prefix, then clustered and numbered by chromosome. The plot is faceted by QTL cluster from left to right, and trait from top to bottom. Within each facet the QTL is on the x-axis and the effect size is on the y-axis. The effect size of the allele is on the scale of the trait (i.e. the marginal change in biomass index or yield in t ha^-1^).

The 4-way MAGIC population used in this study has previously been scored for other traits including root hair length, seedling growth, and traits related to canopy architecture and dormancy (see introduction). Root hair length is of demonstrated value to PUE in both glasshouse and field studies (Gahoonia and Nielsen, 1997, 2003). Delhaize *et al*. (2015) scored two wheat MAGIC populations, including the 4-way population used in this study, for root hair length using rhizosheath size as a surrogate assay. They identified 18 loci linked with this trait. Six of the major QTL mapped to chromosomes 2B, 4D, 5A, 5B, 6A, and 7A and explained between 4.5 and 9.7% of the variation each (Delhaize *et al*., 2015). The QTL identified for root hair length were compared with the QTL in this study. They were clustered in the manner described above and are shown in Figure 7. Five of these six major root hair length loci clustered with PUE QTL (the loci on 5B was the exception). Three other weaker QTL identified in the Delhaize et al. study (there were 18 in total) were also clustered with PUE QTL (Figure 7).

We assume the clusters represent a single gene, and that where the cluster contains multiple QTL it is a single gene influencing multiple traits in multiple environments. Therefore, a cluster would not include more than one QTL for the same trial × trait combination. However, *PUE_4B_3* has two proximate QTL (pue_071 and pue_072, Figure 8) for the same trait and trial, biomass response at Ardlethan 2018, that have founder effects that are different suggesting that there may be two genes in proximity. QTL pue_072 was excluded from subsequent analyses of allele effects as it had smaller effect sizes. Note that the assumption that the clusters represent a single gene can only be confirmed by subsequent sequencing and/or cloning, which is beyond the scope of this study.

The founder effect sizes were used to classify the founder alleles effect as “positive”, “negative”, or “neutral” (the range between the maximum, positive, value and the minimum, negative, value was split into thirds – the middle third being “neutral”). The four traits were then considered in terms of the “productivity” measured, i.e. yield or biomass, and the trait measured, response or resilience.

Within the 56 QTL clusters, if there were pleiotropic QTL where the positive founder for response was also a negative founder for resilience then it was considered to be “antagonistic”. Thirteen of the 56 QTL clusters had an example of pleiotropy of which three clusters had two examples and one had three examples for a total of 18 examples; three for yield and 15 for biomass. All 18 examples had an allele that was antagonistic (i.e. positive for response but negative for resilience). However, because there are four parents to the population there were alternative alleles, and hence 15 of the 18 pleiotropic QTL had alleles that were not antagonistic, including two of the three examples of yield. Note that this approach uses arbitrary assumptions and a very limited definition of “antagonism” and “pleiotropism” to illustrate the extent to which the collection of QTL and clusters is independent with respect to response and resilience.

The effect of combinations of alleles on the four traits in the different trials can be calculated, although the selection of those combinations for the purposes of breeding requires value judgements beyond the scope of this paper (see discussion). For example, the predicted change in trait and trial values for the 6 alleles associated with the greatest root hair length in Delhaize *et al*. (2015) is shown in Figure 9. The change is measured in terms of the percent of the variation attributable to genetics (to normalise for biomass and yield being on different scales: i.e. “lidar index” vs. t ha^-1^). If the effect is due to longer root hairs, then the impact is mixed. For example, it is positive for yield resilience at York in both years and, to a lesser extent, Ardlethan in 2017, and it is also positive for yield response in Ardlethan in 2017. It was negative for response at York 2017, but not 2018, and a negative for yield resilience in Wallaroo.

**Figure 9.**
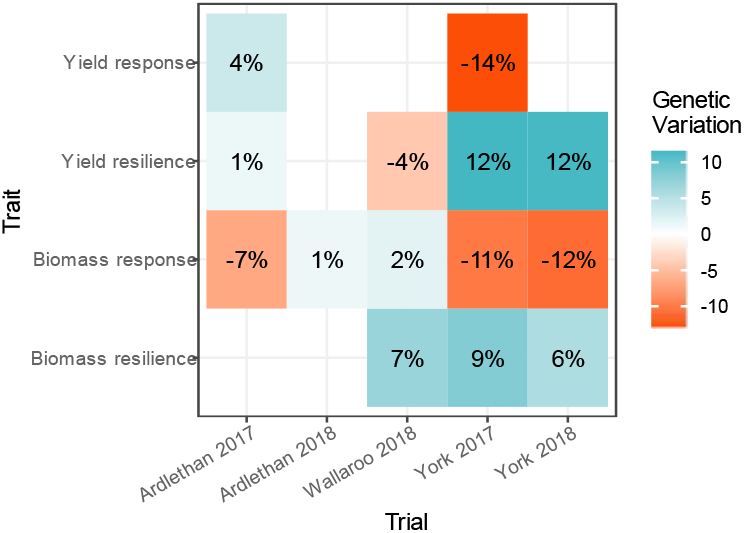
The calculated effect on the variation for traits and trials of the alleles conferring the greatest root hair length in Delhaize et al. (2015). The variation is calculated as the percentage of the trait variation that could be attributed to QTL effects.

## Discussion

### Response and Resilience

This study took an agnostic approach to the mechanisms driving an improvement in PUE. PUE is an aggregate of traits, and from a practical perspective it is not essential to understand what traits underly the QTL. However, by comparing productivity with 0 and 30 kg P ha^-1^ fertilization the response and resilience traits are likely to be associated with distinct physiological mechanisms. For instance, plants that forage the topsoil will be more resilient in soils with low levels of available Pi, whereas plants able to better exploit narrow bands of concentrated P are likely to be the more responsive in fertilized soils.

Response and resilience traits might be antagonistic, particularly from a “rhizoeconomic” perspective (see Lynch *et al*., 2005). An expansive foraging root system may be advantageous under severe deficiency but, where P is abundant, the same trait might represent an unnecessarily high cost to the plant, potentially reducing productivity. The negative correlation of response and resilience in this study is evidence of antagonistic effects of some traits on these two different aspects of PUE; though the trait agnostic approach used does not give insight into which traits are antagonistic. However, there were relatively few examples of antagonistic QTL in this study and they were largely associated with biomass (15 examples) rather than yield (three examples).

One example of antagonistic QTL is associated with root hair length. The cv. Westonia allele associated with cluster PUE_2B_1 was positive for yield resilience at York in 2018 (pue_027) and 2017 (pue_026), but was negative for yield response in 2017 (pue_024, Figure 8). The same allele was also a major positive influence on root hair length in Delhaize *et al*. (2015). This suggests that root hair length had a positive effect on P uptake particularly when foraging, but it is less clear why it would have had a negative impact on yield response and only in 2017. It should be noted that while the effect of *Rhizoctonia* at York in 2017 was scored and accounted for in its effects on yield and biomass if there were any effects that were not associated with the barepatch symptoms they may not have been accounted for. One could speculate that there might be a relationship between root hair length and disease. For example, root hairs are the site of infection for *Pythium* root rot species in wheat (Bruehl, 1953; Royle and Hickman, 1964), although there are no studies examining root hair length and infection density. Furthermore, these alleles had a positive impact on response at Ardlethan in 2017, where there was a substantial deficit in early rainfall and indeed any rainfall. The hairs may have helped take up fertilizer in dry top-soil conditions.

### Achieving the economic optimum

The goal of breeding for enhanced PUE is not to abolish the application of P fertilizers, but rather to reduce the rate of Pi application. Bovill *et al*. (2013) advocate the economic optimum as a goal of PUE breeding: assessing the point at which the marginal cost of applying additional fertilizer meets the marginal increase in profit. As the cost of P fertilizer increases, the economic optimum decreases and therefore the goal of PUE breeding should be to both improve the response to the application of P and improve the ability of the crop to exploit residual soil P. Australian farmers typically band P fertilizers at sowing, a response to the generally P deficient soils. While high fertilizer prices and drought conditions are seeing this practice reduced, it is still likely that fertilizer will be applied to compensate for P removed by the previous year’s crop and at rates that reflect the poor efficiency of uptake. The idiotypic genotype would combine both response and resilience traits; efficiently utilizing a reduced rate of fertilization.

The response of productivity to increasing P fertilization is not linear but a saturating curve. Finding the point of inflection is challenging (Simpson *et al*., 2014, 2015; Haling *et al*., 2016), requiring multiple rates of P application. QTL mapping requires large populations to be meaningful, so treatment numbers must be minimized for trials to be practical. A follow up study could select optimised combinations of response and resilience QTL alleles for particular environments (avoiding and/or compensating for antagonistic effects). Smaller subsets of genotypes in the MAGIC population with these desirable combinations (“tails” similar to the phenotypic approach proposed by Rebetzke *et al*. (2017)) could be assembled and grown at a wider range of intermediate P rates to confirm whether these QTL can be combined to reduce the economic optimum rate of P while maintaining productivity.

### Improving PUE through breeding

Bovill *et al*. (2013) concluded that the lack of a consistent definition of PUE is hampering the genetic improvement of the trait. The response approach of McDonald *et al*. (2015), employed in this study and extended with the concept of resilience, aims to separate the measurement of PUE from absolute productivity.

Nevertheless, response and resilience were positively correlated with productivity with and without fertilizer, respectively. This suggests that response is increasing passively through yield breeding where non-limiting P is provided (as is typically the case). Passive gains in fertilizer use efficiency through yield breeding have been demonstrated (Batten, 1992; Ortiz-Monasterio R. *et al*., 1997). The correlation between 0 kg P ha^-1^ productivity and resilience suggests a breeder with interest in resilience traits could breed for productivity in the absence of P fertilizer. However, it is worth noting that performance in low-input conditions will not necessarily be undermined by breeding and selection in high-input conditions (Voss-Fels *et al*., 2019)

However, as the range of correlations shows, there is also scope for breeding directly for PUE traits. Given (a) the pipeline for the development and release of a new wheat variety can be 5-10 years and (b) long term fertilizer price increases can be predicted with some certainty it may be profitable to breed specifically for improved PUE. A QTL study, like this one, also allows for the development of marker assisted selection approaches within breeding programs. Breeders are adopting genomic selection strategies, as sequencing and mapping costs drop. PUE QTL can help to define the breeding values for a marker matrix, developing a variety optimized not for current fertilizer prices, but the prices expected 10 years hence when the variety comes to market. Trait relevant markers have been shown to increase prediction accuracy when paired with genomic selection in maize (Liu *et al*., 2019).

### Future directions and conclusion

Improving crop PUE is an intrinsically valuable goal considering the increasing cost and importance of P as agricultural input. However, a breeding program focused on PUE will have to compete with other priorities such as disease resistance and adaptation to climate change. The identification of QTL allows the development of trait relevant markers that can be integrated into marker assisted selection or genomic selection approaches. Trialling “tail” combinations of these QTL at a wider range of fertilization rates could help to establish the effect on the economic optimum P fertilization rate.

## Authors’ Contributions

PRR and ED conceived of the study. The experiments were designed and executed by APW, GP, and CW. The layouts were designed by ABZ and APW. The data was analysed by ABZ, APV, and APW. APW wrote the manuscript with input from ABZ, APV, PRR and ED.

## Acknowledgements

We are profoundly grateful to farmers Grant Kitto and Charlie Boyle for hosting our trials. We thank John Byrne, Tom McLucas, Savannah McGuirk, and Geetha Perera for their dedicated efforts in maintaining, sampling, and processing the experiments. We thank Jamie Scarrow and Michael Salim of the Australian Plant Phenomics Facility for excellent support with the Phenomobile Lite. This work was supported by a Grains Research and Development Corporation grant CFF00009 *Molecular markers for root hair traits and enhanced phosphorus use efficiency (PUE) in wheat*.

## Abbreviations

BLUPs: Best Linear Unbiased Predictors;
DH: Doubled Haploid;
Pi: Inorganic Phosphate;
LIDAR: Light Detection and Ranging;
MET: Multi-Environment Trial;
MAGIC: Multiparent Advanced Generation InterCross;
P: hosphorus;
PUE: P use efficiency;
QTL: Quantitative Trait Loci;
RIL: Recombinant Inbred Line;
WGAIM: Whole Genome Average Interval Mapping

## Supplementary Information

**Supplementary Figure 1.**
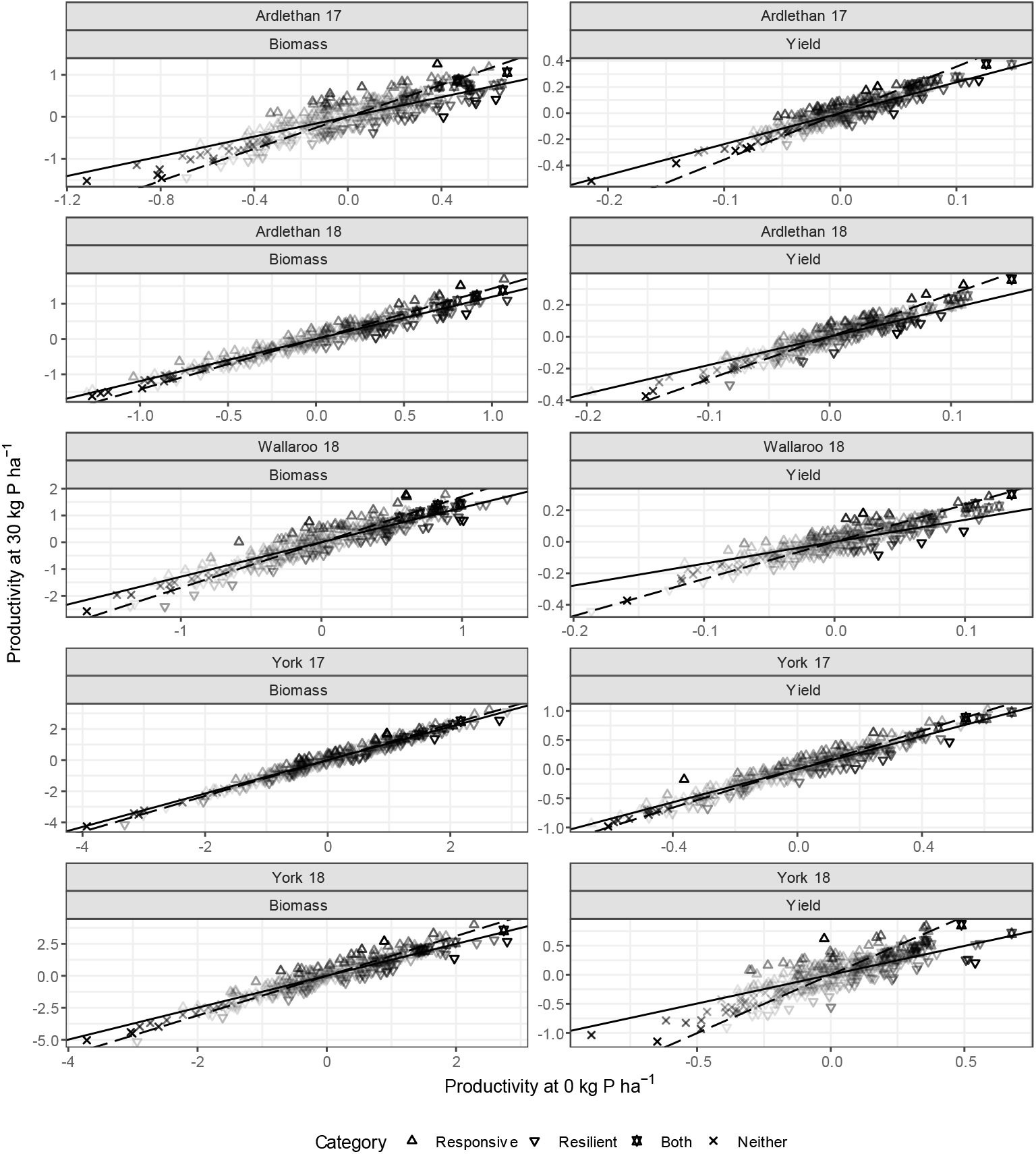
An expansion of Figure 4 to include all trials (vertical facets) and productivities (horizonal facets). Yield at 0 kg P ha^-1^ is plotted against yield at 30 kg P ha^-1^. Lines for the average genetic yield response to fertilisation (**solid**) and average genetic yield resilience in the absence of fertilization (**dashed**) in this trial. The difference in response for each genotype from the population average is its vertical distance (y-axis) from the average, i.e. the red line. The difference in resilience for each genotype from the population average is its horizontal distance (x-axis) from the average, i.e. the blue line. Each genotype has been categorised and coded by shape. If it is **responsive**, i.e. above the solid line, it is a **triangle**; if it is **resilient**, i.e. right of the dashed line, it is an **inverted triangle**; if it is **both** resilient and responsive, i.e. above the solid line and right of the dashed line, it is a **star**; and if it is **neither** resilient nor responsive, i.e. below the solid line and left of the dashed line, it is a **cross**. The shading of the point indicates the degree of response and/or resilience, and highlights that the measure is independent of yield at either P level. For the genotypes that are neither or both responsive and resilient the shading is the average of the two measures and will increase with distance from the origin. The slope of the regression is derived from a factor analytic model fitted across all five trials in the study and represents the genetic component of the response; it does not reflect a regression of the BLUPs for the two P fertilization levels.

**Supplementary Figure 2.**
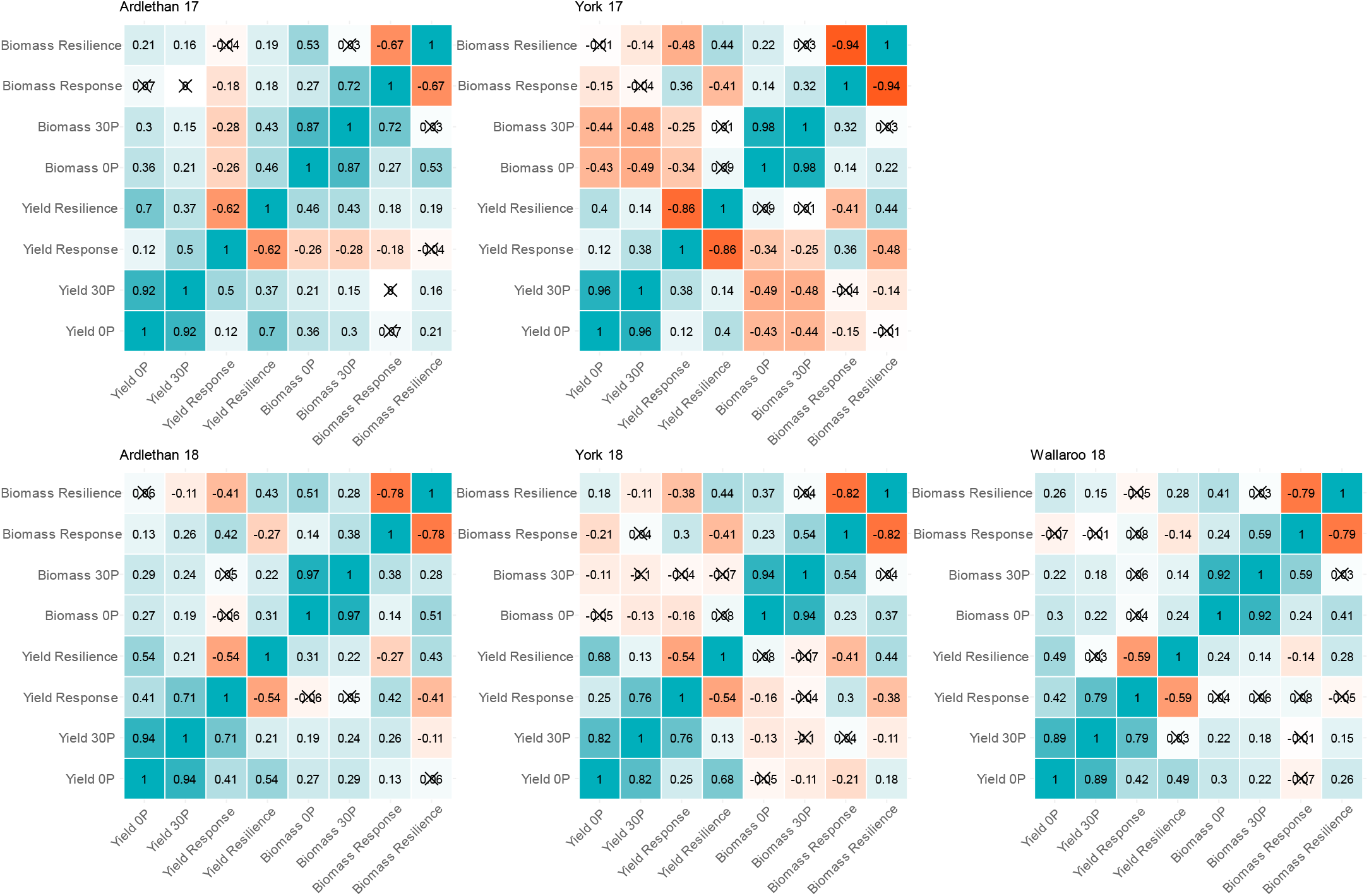
Correlograms for each BLUP faceted by each trial. Each tile is coloured and labelled by the size and direction of the correlation (Pearson correlation coefficient). Correlations marked with a cross are insignificant.

